# A mechanistic switch from selective transporter to an ion channel impairs the filamentation signalling capability of ammonium transceptors in yeast

**DOI:** 10.1101/2021.08.25.457613

**Authors:** Gordon Williamson, Ana Sofia Brito, Adriana Bizior, Giulia Tamburrino, Gaëtan Dias Mirandela, Thomas Harris, Paul A. Hoskisson, Ulrich Zachariae, Anna Maria Marini, Mélanie Boeckstaens, Arnaud Javelle

## Abstract

Ammonium translocation through biological membranes by the ubiquitous Amt-Mep-Rh family of transporters plays a key role in all domains of life. Two highly conserved histidine residues protrude into the lumen of these transporters, forming the family’s characteristic Twin-His motif. It has been hypothesized that the motif is essential to confer the selectivity of the transport mechanism. Here, using a combination of *in vitro* electrophysiology, *in vivo* yeast functional complementation and *in silico* molecular dynamics simulations, we demonstrate that variations in the Twin-His motif trigger a mechanistic switch between a specific transporter, depending on ammonium deprotonation, to an unspecific ion channel activity. We therefore propose that there is no selective filter that governs the specificity in Amt-Mep transporters but the inherent mechanism of translocation, dependent on the fragmentation of the substrate, ensures the high specificity of the translocation. We further show that both mechanisms coexist in fungal Mep2 Twin-His variants, disrupting the transceptor function and so inhibiting the filamentation process. These data strongly support a transport mechanism-mediated signalling process in the long-standing debate on the sensory function of Mep2-like transporters.

## Introduction

Transport of ammonium across biological membranes is a highly conserved and fundamental biological process. Ammonium represents a major nitrogen source for higher plants and is a preferred nitrogen source for many bacterial and yeast species, including *Escherichia coli* and *Saccharomyces cerevisiae* ^*1*^.In humans, ammonium plays an important role in systemic pH control while it is also a cytotoxic metabolic waste product whose accumulation can lead to lethal encephalopathy and pathophysiology ^2,3^.

Cellular ammonium transport is facilitated by the ubiquitous Amt-Mep-Rh superfamily; members of which have been identified in every branch of the tree of life ^4^. The Amt-Mep ammonium transporters are found in bacteria, fungi, plants and invertebrates ^5,6^. Rhesus (Rh) proteins, whilst originally of interest for their central role in the blood group complex, are the mammalian equivalents of the Amt-Mep ammonium transporters ^7,8^.

The physiological relevance of Amt-Mep proteins extends beyond their role in ammonium acquisition as a nitrogen source. In fungi for instance, in the presence of very low ammonium concentrations, specific Amt-Mep transporters, the Mep2-like proteins, have been proposed to act as sensors required for the development of filamentous growth, a dimorphic switch associated to the virulence of pathogenic fungi ^9^. The baker yeast *S. cerevisiae* possesses three Amt-Mep proteins (Mep1-3) and only *mep2Δ* cells are unable to filament, *MEP1* or *MEP3* deletion having no impact on this dimorphic switch. Single Mep2 expression is further sufficient to trigger filamentation and this protein has therefore been suggested to act as a transceptor, able to combine a function of transport and receptor coupled to a signal transmission.

Despite the relevance of this protein family, the transport mechanism of Amt-Mep-Rh has remained largely elusive for decades. Crystal structures of various family members revealed a trimeric organisation with a narrow conducting transport pore through each monomer lined with hydrophobic residues ^10-15^. The hydrophobicity of the central pore represents a high energetic barrier to the movement of ions, and thus it was concluded that Amt-Mep-Rh translocate the neutral form NH_3_ ^11^. While it was next shown that Amt-Mep-Rh proteins mediate NH_4_^+^ deprotonation ^16^, functional studies demonstrated that *Archaeoglobus fulgidus* Amt1 and 3 and *E. coli* AmtB mediate electrogenic ammonium transport ^17-19^

AmtB (the native *E. coli* ammonium transporter) is the paradigmatic member of the Amt-Mep-Rh family, and is widely used as a model system to study ammonium uptake ^17,20,21^. Recently, we reported a novel mechanism for ammonium-mediated transport in *E. coli* AmtB ^22^. Two highly conserved histidine residues, H168 and H318, protrude into the lumen of the pore, forming the family’s characteristic Twin-His motif ^22,23^. We identified two water wires, connecting the periplasmic side to the cytoplasmic vestibule of AmtB. These two water wires are interconnected via the H168 residue of the Twin-His motif which acts as a seal, preventing the formation of a continuous water chain from the periplasm to the cytoplasm. We showed that, after deprotonation of NH_4_^+^ at the periplasmic side, the two interconnected water wires and the Twin-His motif enable H^+^ transfer into the cytoplasm. A parallel pathway, lined by hydrophobic groups within the protein core, facilitates the simultaneous transfer of uncharged NH_3_. Thus, the Twin-His motif appears essential to the transport mechanism. However, despite its high level of conservation, the Twin-His motif is not universally conserved ^23,24^. For instance, a specific substitution evolved and is tolerated in some fungal Amt-Mep proteins. Specifically, the first histidine is present in the transceptor Mep2-type proteins, whereas the fungal Mep1- and Mep3-type proteins feature a natural occupation by a glutamic acid at the first histidine position, defining two functional Mep-Amt subfamilies in fungi. More recently, functional characterization in *Xenopus* oocytes revealed that *S. cerevisiae* Mep2 mediates electroneutral substrate translocation while Mep1 performs electrogenic transport ^25^. We therefore sought to analyse the role of the Twin-His motif on the transport mechanism of Mep-Amt proteins and the impact of its modification on the ability of fungal Mep2 transceptors to allow filamentation.

Here, we report that altering the Twin-His motif within the pore of *E. coli* AmtB, *S. cerevisiae* and *C. albicans* Mep2 does not impair ammonium transport activity but abolished the pore selectivity against competing monovalent cations. We further demonstrate that this loss of selectivity is the result of a mechanistic switch from a transporter-like activity to a channel-like activity governed by a change in the hydrophobicity of the pore. Finally, we show that this mechanistic switch impacts on the ability of fungal Mep2 proteins to act as transceptors in the development of pseudohyphal growth. These findings show that the mechanism of substrate transport ensures the high specificity of the transport and strongly support the hypothesis that the mechanism of transport of Mep2-like proteins is responsible for the signal that leads to yeast filamentation.

## Results

### Altering pore hydrophobicity does not disrupt transport activity of AmtB

To determine the effect of altering the hydrophobicity of the pore on transport activity, we first looked at the effect of single acidic substitutions H168E and H168D within the Twin-His motif of AmtB, mimicking the substitution present in the fungal Mep1/3-type of transporters. We purified and reconstituted both variants into liposomes and measured their activities *in vitro* using Solid Supported Membrane Electrophysiology (SSME) and *in vivo* by yeast complementation assays. In proteoliposomes containing AmtB^H168D^ or AmtB^H168E^, an ammonium pulse of 200 mM elicited very high amplitude currents of 14.23 nA and 22.28 nA respectively, compared to 3.38 nA observed for the WT (Figure 1A). Importantly, the lifetime of the currents measured for both variants was dependent on liposomal Lipid:Protein Ratio (LPR), indicating that the currents account for a full translocation cycle (Table 1). Additionally, we measured an increase of the catalytic constants (*K*_*m*_) for both variants compared to the WT (Figure 1C, Table 2).

**Figure 1:**
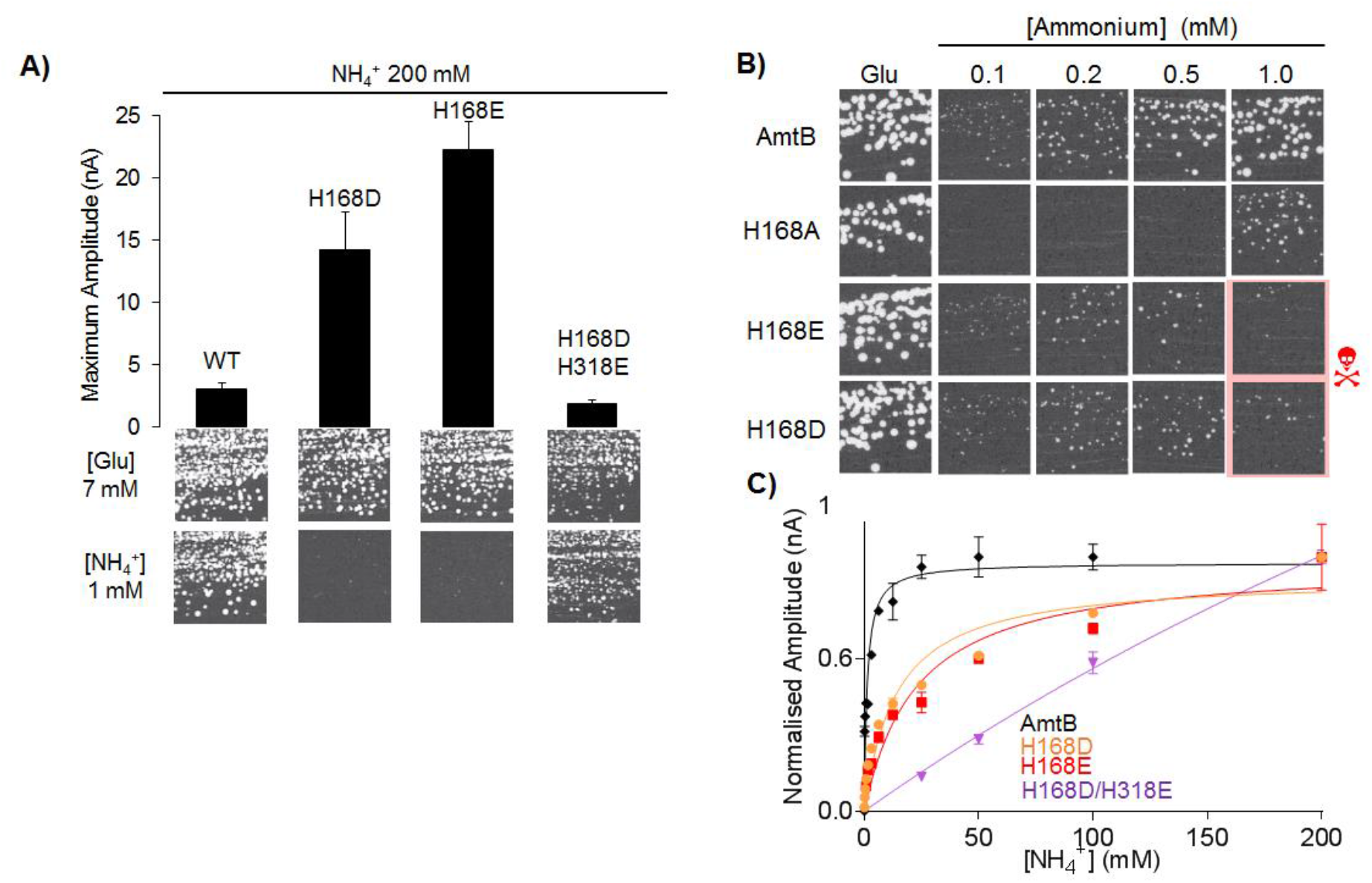
Effect of Twin-His Substitution on AmtB ammonium transport Activity. **A)** *Upper panel*: maximum amplitude of the transient currents measured using SSME following a 200 mM ammonium pulse. *Lower panel*: yeast complementation test after 3 days at 29°C, on glutamate (Glu, positive growth control) or ammonium as a sole nitrogen source. The strain 31019b (*mep1*Δ *mep2*Δ *mep3*Δ *ura3*) was transformed with the pDR195 plasmid allowing expression of the various AmtB variants. **B)** Yeast complementation test, after 5 days at 29°C, in the presence of a range of ammonium concentrations (0.1 mM – 1.0 mM) or glutamate. **C)** Kinetics analysis for the transport of ammonium using SSME. The maximum amplitudes recorded after a 200 mM ammonium pulse have been normalised to 1.0 for comparison.

**Table 1:** Decay time constants (s^−1^) of transient currents triggered after an ammonium, methylammonium or potassium pulse of 200 mM in proteoliposomes containing AmtB variants at various LPR*.

| Variant | Ammonium |  | MeA |  | K<sup>+</sup> |  |
| --- | --- | --- | --- | --- | --- | --- |
|  | LPR 10 | LPR 5 | LPR 10 | LPR 5 | LPR 10 | LPR 5 |
| WT | 13.4 ± 1.5 | 18.7 ± 1.0 | 3.6 ± 0.5 | 8.3 ± 1.2 | NC | NC |
| H168D | 31.4 ± 1.3 | 38.1 ± 4.1 | 17.7 ± 2.1 | 26.4 ± 4.2 | NC | NC |
| H168E | 45.5 ± 4.8 | 53.9 ± 1.6 | 27.0 ± 1.9 | 36.4 ± 3.1 | NC | NC |
| H168D/H318E | 15.8 ± 1.1 | 17.8 ± 0.7 | 10.5 ± 2.5 | 13.9 ± 1.8 | 2.3 ± 0.4 | 4.9 ± 0.2 |

\*NC: No transient current recorded

**Table 2:** *Km* (mM) of AmtB variants for ammonium, methylammonium, and potassium, using SSME. *

| Variant | NH<sub>4</sub><sup>+</sup> | MeA | K<sup>+</sup> |
| --- | --- | --- | --- |
| WT | 0.8 ± 0.1 | 49.86 ± 4.76 | NC |
| H168D | 14.02 ± 4.08 | 55.09 ± 12.67 | NC |
| H168E | 12.87 ± 7.79 | 86.00 ± 9.46 | NC |
| H168D/H318E | NA | NA | NA |

\*NC: No transient current recorded. NA: Not Applicable

Expressed in a *S. cerevisiae* triple-*mepΔ* strain, deprived of its three endogenous Mep ammonium transporters, both variants were unable to restore growth on 1 mM ammonium after 3 days, suggesting a loss of function (Figure 1A). However, after 5 days, an inhomogeneous weak growth was observed (Figure 1B). Previous work has demonstrated that non-controlled ammonium influx can be toxic to *S. cerevisiae* ^26,27^. To test if this inhomogeneous growth could be linked to toxic ammonium influx, the yeast complementation assay was repeated with a lower range of ammonium concentrations (0.1-1.0 mM) (Figure 1B). The complementation ensured by both AmtB^H168D^ and AmtB^H168E^ variants was improved by decreasing the ammonium concentrations. These results suggest that the acidic substitution at position 168 drastically increases the ammonium flux through AmtB, rendering it toxic to yeast cells. We did not previously observe ammonium toxicity with the AmtB^H168E^ variant expressed in yeast ^24^. In these experiments, AmtB^H168E^ was expressed under the control of the *MET25* promoter and in the presence of ammonium sulfate, which may partially repress the *MET25* promoter. In the present experiment, the variants are expressed under the control of the *PMA1* promoter, a stronger promoter than the one of *MET25*. Therefore, enhanced expression of the variants in the present experiment could explain the ammonium toxicity observed.

To further probe the impact of pore hydrophobicity on the activity and specificity of AmtB, we substituted both residues of the Twin-His motif with acidic residues (AmtB^H168D/H318E^). SSME measurements revealed that a 200 mM NH_4_^+^ pulse elicited an LPR-dependent transient current with a maximum amplitude of 1.84 nA in proteoliposomes containing AmtB^H168D/H318E^, a 1.8-fold reduction compared to WT AmtB (Figure 1A, Table 1). Additionally, a catalytic constant (*K*_*m*_) for AmtB^H168D/H318E^ could not be determined as saturation could not be achieved, even following an ammonium pulse of 200 mM (Figure 1C, Table 2). When expressed in the *S. cerevisiae* triple-*mepΔ* strain, AmtB^H168D/H318E^ was able to restore cell growth on low ammonium concentrations (Figure 1A). These data demonstrate that, while AmtB^H168D/H318E^ is still functionally active, its activity seems to be reduced compared to WT AmtB. The absence of saturable kinetics further suggests that the variant behaves like a channel rather than having transporter-like activity. We have previously observed this type of behaviour when we replaced the Twin-His motif by alanine residues although the mechanism of this switch remained elusive ^22^.

Previously, methylammonium (MeA) was used as a substrate analogue for ammonium. However, it triggers transient currents of only 15-20% of those elicited by ammonium, indicating strong substrate discrimination by AmtB ^19,20,28^. To determine if this level of discrimination is maintained in the Twin-His variants, the AmtB^H168D^ and AmtB^H168E^ variants were subjected to a pulse of 200 mM MeA during SSME. Both AmtB^H168D^ and AmtB^H168E^ exhibited increased activity compared to WT AmtB (9-fold and 12-fold increase in current amplitude respectively). The double mutant AmtB^H168D/H318E^ shows a maximum amplitude comparable to WT AmtB (Figure 2A) but it was not possible to determine a catalytic constant, again due to the lack of saturation (Figure 2B, Table 2). These data show that the mutations do not alter the ability of AmtB to discriminate between MeA and ammonium *in vitro*. Yeast complementation was carried out in parallel. MeA cannot be metabolized by yeast cells and is toxic at high external concentrations ^24,29^. The native AmtB and all the Twin-His variants allowed growth of triple-*mepΔ S. cerevisiae* cells on glutamate media, but not in the medium supplemented with MeA (Figure 2A). This confirms that all the variants are active in transporting MeA in yeast.

**Figure 2:**
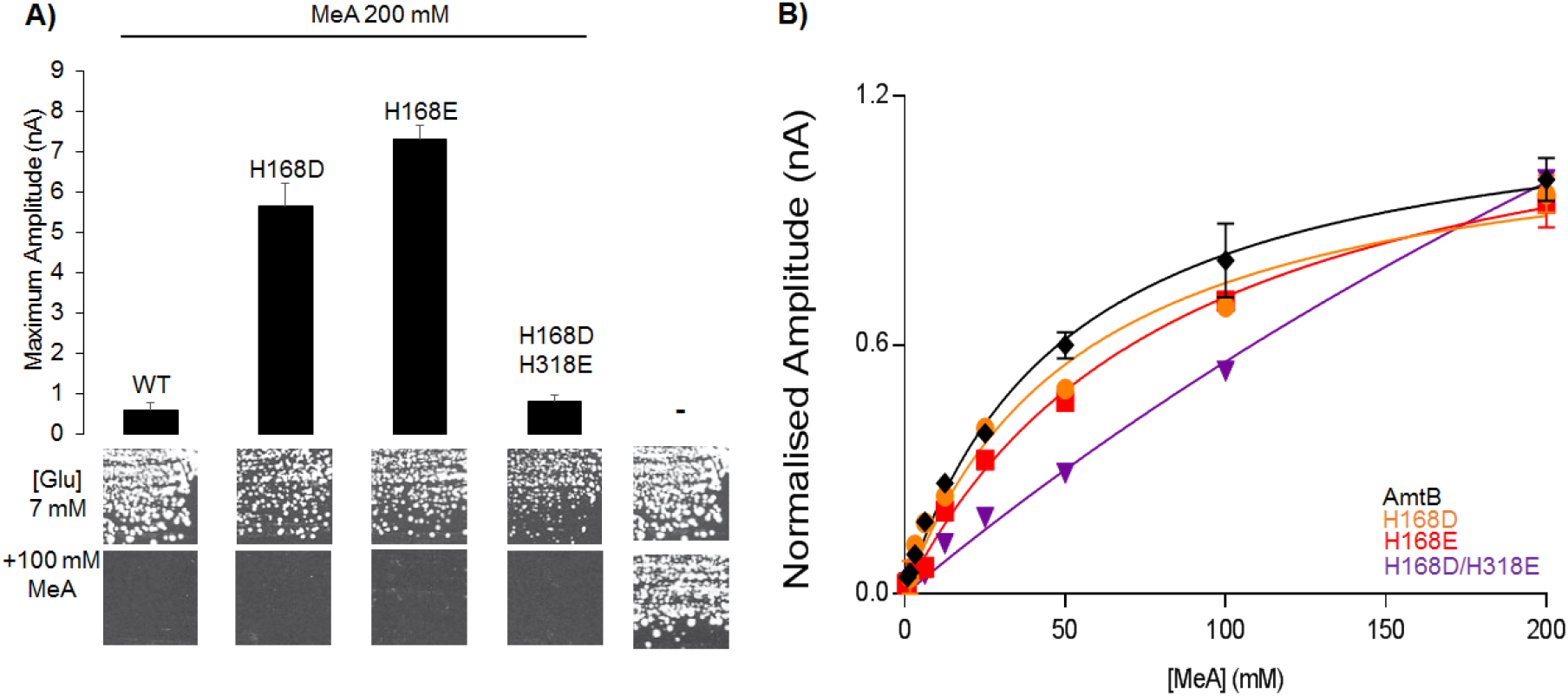
Effect of Twin-His Substitution on AmtB methylammonium transport Activity. **A)** *Upper panel*: maximum amplitude of the transient current measured using SSME following a 200 mM methylamine (MeA) pulse. *Lower panel*: yeast complementation test, after 3 days at 29°C, on solid minimal medium containing, as the sole nitrogen source, glutamate 7 mM (glu, positive growth control) supplemented or not with 100 mM methylammonium (MeA). The strain 31019b (*mep1Δ mep2Δ mep3Δ ura3*) was transformed with the empty pDR195 plasmid (-), or with pDR195 allowing expression of the various AmtB variants. **B)** Kinetics analysis for the transport of methylammonium (MeA) using SSME. The maximum amplitudes recorded after a 200 mM MeA pulse have been normalised to 1.0 for comparison.

### Altering pore hydrophobicity abolishes transport selectivity

The findings reported above showed that altering the hydrophobicity of the pore does not inactivate AmtB but had a substantial impact on the transport mechanism. Thus, we next investigated whether the substitutions also affect the ammonium/Mea selectivity of AmtB against competing ions.

A 200 mM K^+^ pulse failed to elicit a measurable current in AmtB^H168D^ or AmtB^H168E^ proteoliposomes, but was able to trigger a clear transient current in AmtB^H168D/H318E^ (Figure 3A). The amplitude and decay time of the currents recorded for AmtB^H168D/H318E^ are LPR-dependent, confirming that they are caused by K^+^ translocation and not only the result of protein-substrate interaction (Figure 3B, Table 1). It was not possible to determine a catalytic constant for AmtB^H168D/H318E^, again due to the lack of saturation, indicating a channel-like rather than a transporter-like activity (Figure 3C, Table 2). Moreover, all the variants but not the WT were able to complement the growth defect of a triple-*mepΔ trkΔ S. cerevisiae* strain, which lacks the three endogenous ammonium (Mep) transporters and the 2 major potassium (Trk) transporters, in the presence of a limited concentration of K^+^ (Figure 3A). The complementation is clearly improved when AmtB^H168D/H318E^ is expressed compared to AmtB^H168D^ or AmtB^H168E^, showing that the single variants are also able to translocate potassium albeit at a lower rate than AmtB^H168D/H318E^. This could explain why we measured a current after a K^+^ pulse in the proteoliposomes containing AmtB^H168D/H318E^ but not with AmtB^H168D^ or AmtB^H168E^. We reason that there could be an inverse relationship between the hydrophobicity of the central pore and the selectivity of AmtB.

**Figure 3:**
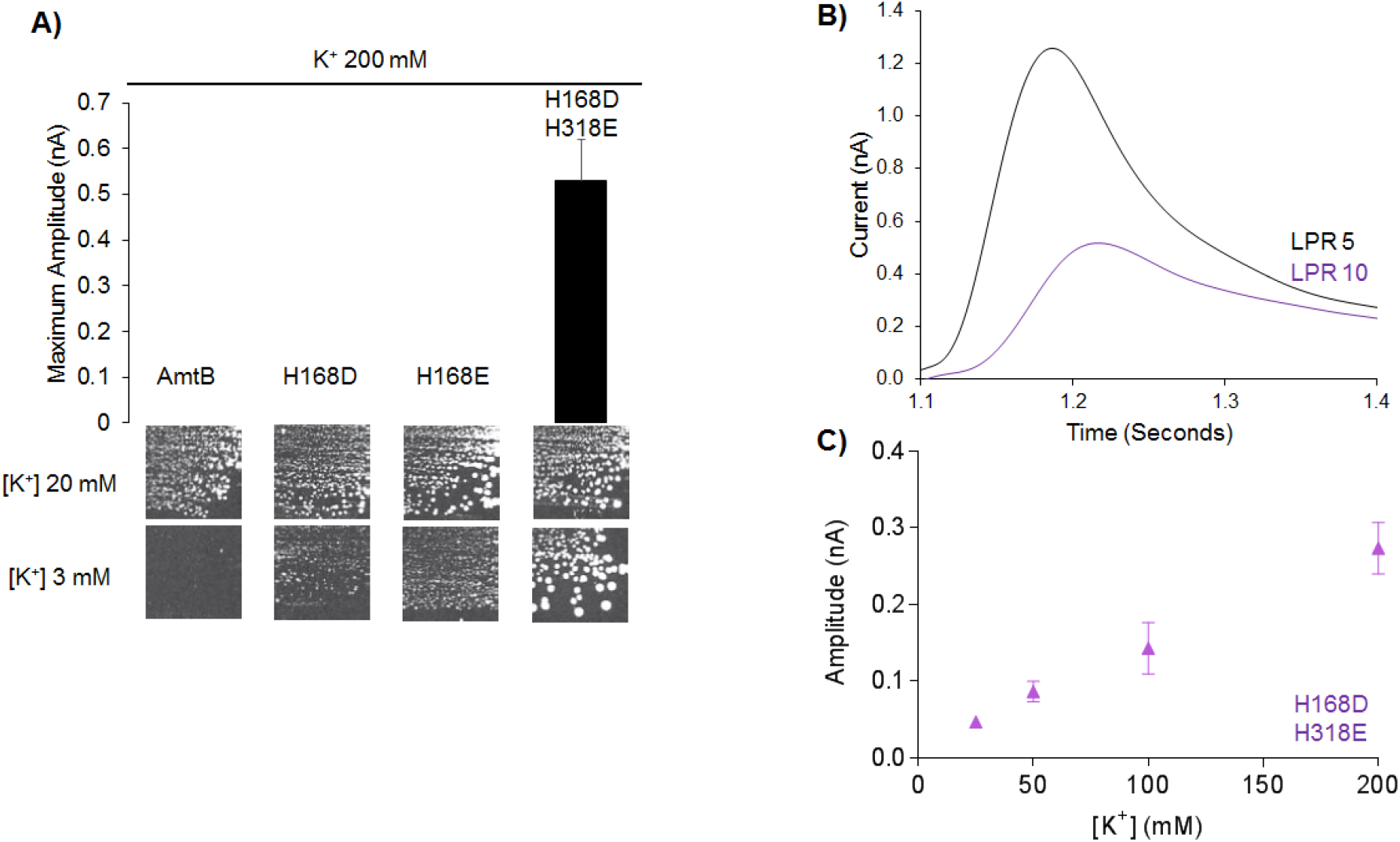
K^+^ transport in AmtB Twin-His Variants. **A)** *Upper panel*: maximum amplitude of the transient current measured using SSME following a 200 mM potassium pulse. *Lower panel*: yeast complementation test, after 5 days at 29°C, in the presence of 20 mM or 3 mM potassium. The #228 strain, *mep1Δ mep2Δ mep3Δ trk1Δ trk2Δ leu2*, was transformed with the pDR195 plasmid allowing expression of the various AmtB variants. **B)** Transient currents measured using SSME following a 200 mM ammonium pulse with AmtB^H168D/H318E^ reconstituted into proteoliposomes at a LPR 5 (black) or 10 (purple). **C)** Kinetics analysis for the transport of potassium using SSME.

### Loss of AmtB selectivity is due to increased pore hydration accompanied by a mechanistic change

To understand the loss of substrate selectivity observed in the AmtB variants and the switch from transporter-like to channel-like activity observed in the SSME recordings with the AmtB^H168D/H318E^ variant, molecular dynamics simulations were conducted. Our previously proposed model for the transport mechanism of AmtB suggests that, after deprotonation of NH_4_^+^ at the periplasmic side, two interconnected water wires enable H^+^ transfer into the cytoplasm. A parallel pathway, lined with hydrophobic groups within the protein core, facilitates the simultaneous transfer of NH_3_ (Figure 4A).

**Figure 4:**
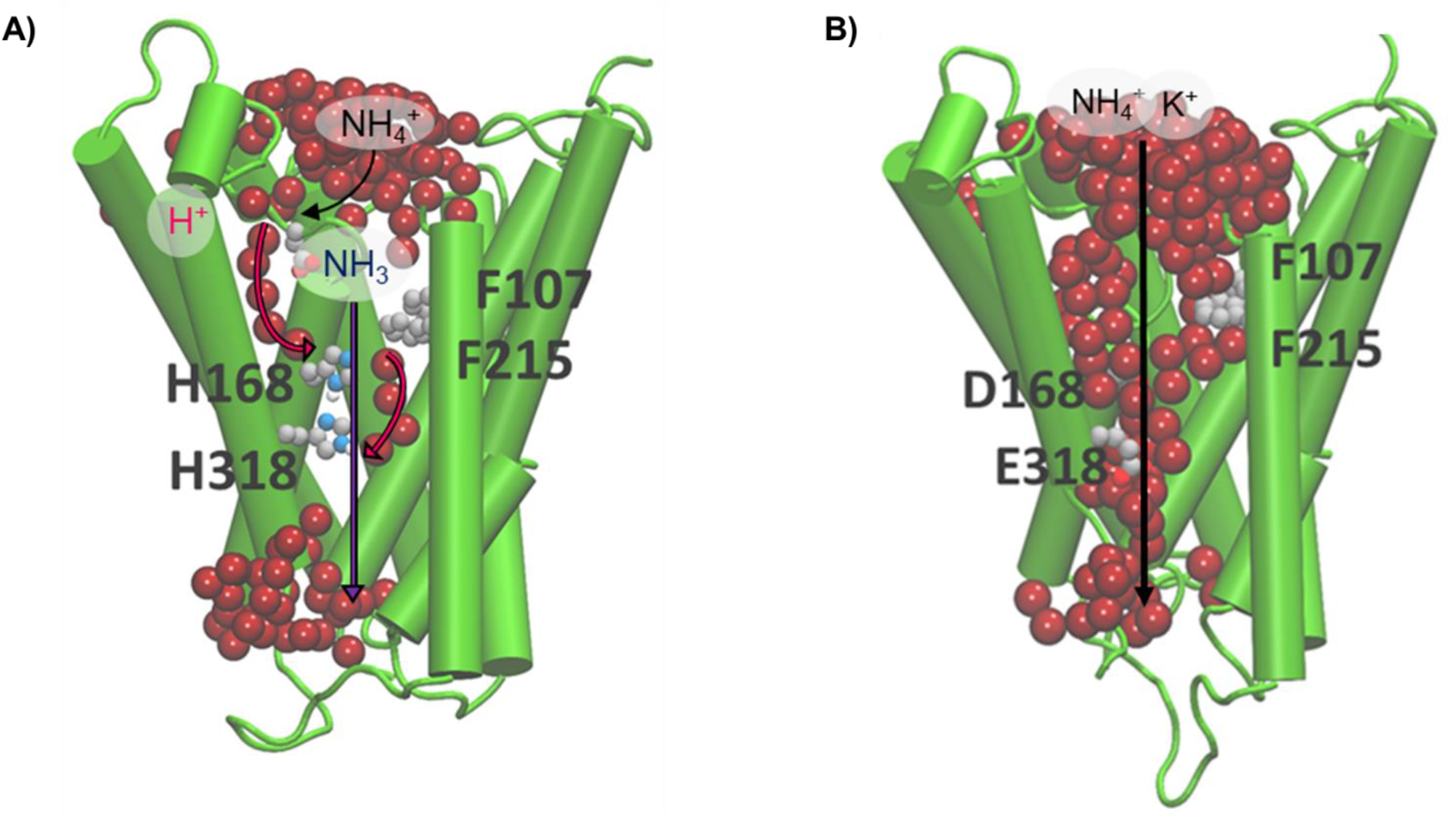
Schematic Comparison of Transport in WT AmtB and AmtB^H168D/H318E^. **A)** Molecular dynamic simulation of AmtB monomer, showing the interconnected water wires (represented as red spheres). Following sequestration of NH_4_^+^ at the periplasmic face, NH_4_^+^ is deprotonated and H^+^ and NH_3_ follow two separate pathways to join the cytoplasm (magenta arrows depict the pathway for H^+^ transfer, dark blue arrows for NH_3_) facilitated by the presence of two internal water wires. **B)** Molecular dynamic simulation of AmtB^H168D/H318E^ monomer, showing the pore filled with water molecules. Due to the increased hydration within the pore, periplasmic NH_4_^+^ and K^+^ are translocated directly through the central pore to the cytoplasm.

Molecular dynamics Simulations based on alterations to the Twin-His motif through the introduction of charged residues show that these destabilise and widen the pore, causing it to fill with water and forming a continuous aqueous channel (Figure 4B). This represents a significant change from the ordered single-file chains of water molecules, separated by the Twin-His motif, observed in the simulations with the WT. The formation of a wide aqueous pore at the expense of discrete water chains disrupts the mechanism of transport and compromises the ability of AmtB to act as a specific transporter, since the newly flooded pore can enable the direct passage of hydrated cations.

To test if the single-file water wires are indeed disrupted by the substitutions, we employed a D_2_O-based assay. Because the strength of a covalent bond involving deuterium increase compared to hydrogen, proton mobility is reduced by 30% for each D_2_O molecule compared to H_2_O ^30^. If the water wires are intact and are required for the transport mechanism, replacement of H_2_O with D_*2*_O is expected to result in the complete abolishment of current when measured by SSME. If, however, the mechanism does not require the water wires, the substitution of H_2_O by D_2_O should not affect the observed current following an ammonium pulse.

As previously observed in WT AmtB, no current was measured in D_2_O conditions (Figure 5) ^22^. In AmtB^H168D^ and AmtB^H168E^, the current observed following a 200 mM ammonium pulse was diminished by about 4-fold in the presence of D_2_O compared to H_2_O, but not completely abolished (Figure 5). The stark reduction of current in the presence of D_2_O indicates that proton hopping remains a mechanistic feature within AmtB^H168D^ and AmtB^H168E^. However, the lack of complete abolition implies that some charge translocation is occurring by another mechanism. These data suggest that two transport mechanisms are used by these variants, one depending on NH_4_^+^ deprotonation, and the other allowing the occasional passage of hydrated NH_4_^+^ or K^+^, as shown in Figure 4. This result is in agreement with the capacity of these variants to transport K^+^ in yeast (Figure 3A).

**Figure 5:**
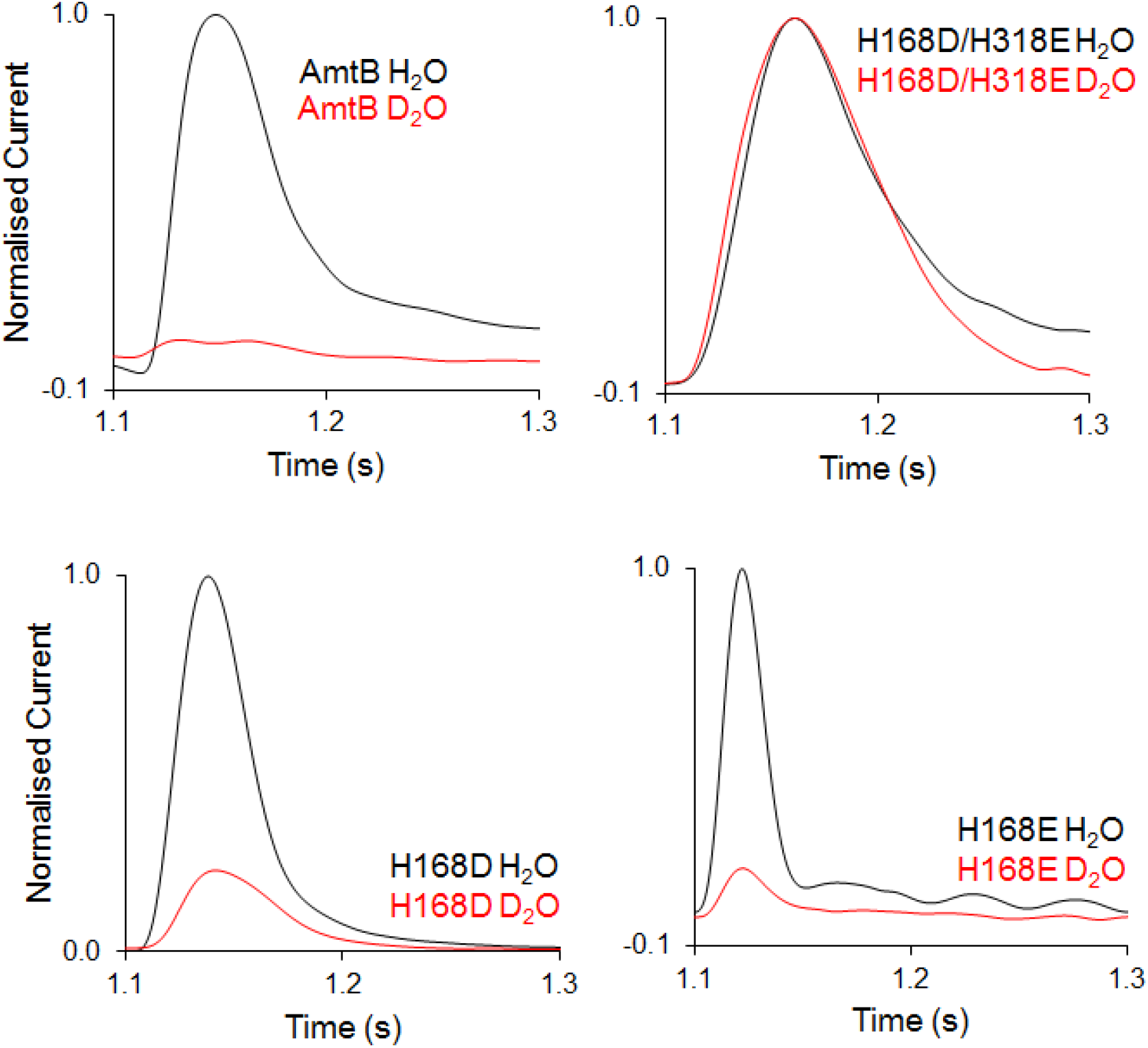
Hydrophobicity of AmtB transport pore Governs Mechanistic Switch. Transient currents measured using SSME following a 200 mM ammonium pulse on sensors prepared with solutions containing either H_2_O (black) or D_2_O (red) in WT AmtB, AmtB^H168D/H318E^, AmtB^H168D^ or AmtB^H318E^. D_2_O currents have been normalised to respective H_2_O currents.

In contrast, in AmtB^H168D/H318E^, a 200 mM ammonium pulse elicits transient currents of similar magnitude with either D_2_O or H_2_O (Figure 5). This indicates that the central mechanism of proton hopping in WT AmtB is no longer a mechanistic feature of AmtB^H168D/H318E^ which become able to directly transport NH_4_^+^ without deprotonation. These findings demonstrate that hydrophilic substitutions within the Twin-His motif gradually lead to a switch in the transport mechanism of AmtB from transporter to channel-like activity, which is fully observed for AmtB^H168D/H318E^. In the latter, NH_4_^+^ is translocated as an intact cation in its hydrated form, abolishing transport specificity (Figure 4). These data also explain why the ammonium/MeA/potassium transport activity cannot be saturated in proteoliposomes containing AmtB^H168D/H318E^ (Figure 1-3, Table 2).

### The Twin-His motif is involved in the transport specificity of ScMep2

Previously, we showed that alanine mutations in the Twin-His motif (AmtB^H168A^ and AmtB^H318A^) also alter the selectivity of the AmtB pore, resulting in the ability of the variants to translocate K^+^ ions^22^. To understand the general importance of the hydrophobicity of the central pore in substrate selectivity, we mutated the Twin-His residues of the *S. cerevisiae* Mep2 into alanine or glutamate residues. We tested if the *S. cerevisiae* triple-*mep*Δ strain expressing ScMep2^H194A^, ScMep2^H194E^, ScMep2^H348A^, or ScMep2^H348E^ was able to grow on low ammonium, similarly to cells expressing native ScMep2 (Figure 6A). We found that a single substitution of the first or second histidine of the Twin-His motif into alanine or glutamate does not affect ammonium transport function of ScMep2. However, contrary to what is observed with AmtB, the ScMep2 double Twin-His mutants (ScMep2^H194A/H348A^ and ScMep2^H194E/H348E^) do not enable growth of the triple-*mep*Δ cells on low ammonium (Figure 6A). We also tested the capacity of ScMep2 variants to intoxicate cells in the presence of high MeA concentrations. Contrary to ScMep1, native ScMep2 is unable to intoxicate cells in the presence of MeA, proposed to be due to a lower maximal transport rate ^31^. Expression of ScMep2 variants with single His substitutions reduces growth of the triple-*mep*Δ cells in the presence of MeA, suggesting a higher transport flux through the variants or an altered transport mechanism increasing the sensitivity of the cells to the toxic compound (Figure 6A). The substitution of both histidines in the Twin-His motif is not accompanied by increased sensitivity of the cells to Mea, which is in agreement with the absence of ammonium transport by the ScMep2^H194A/H348A^ and ScMep2^H194E/H348E^ variants (Figure 6A). As for AmtB, the Twin-His substitutions of ScMep2 do not affect ammonium/MeA discrimination.

**Figure 6:**
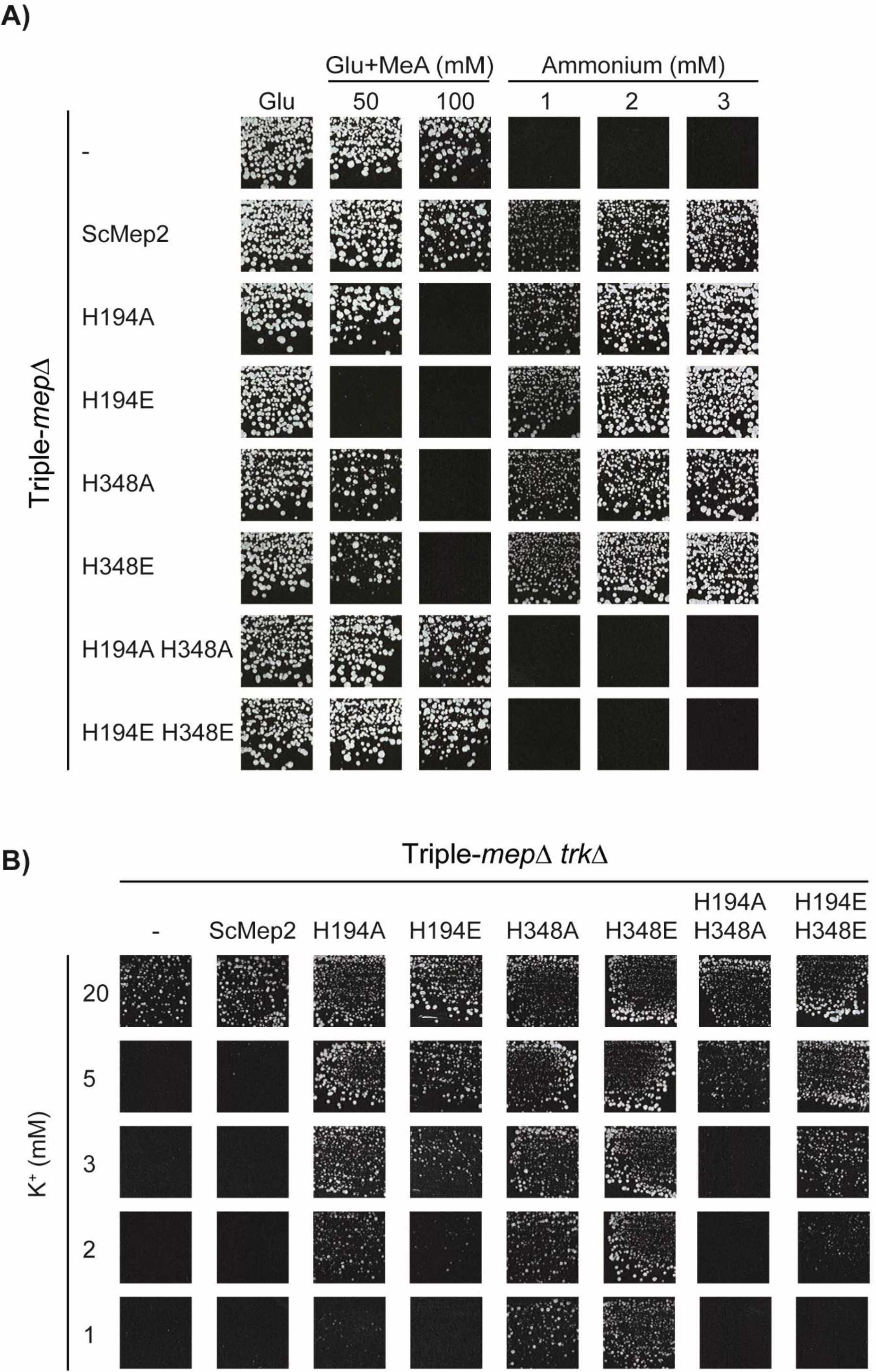
Effect of Twin-His substitutions on ScMep2 transport activity and specificity. **A)** Growth tests, after 4 days at 29°C, on solid minimal medium containing, as the sole nitrogen source, ammonium 1, 2 and 3 mM or potassium glutamate (Glu, positive growth control) supplemented or not with 50 or 100 mM methylammonium (MeA). The *Saccharomyces cerevisiae* strain 31019b (*mep1Δ mep2Δ mep3Δ ura3*) was transformed with the pFL38 empty plasmid (-) or with YCpMep2, YCpMep2^H194A^, YCpMep2^H194E^, YCpMep2^H348A^, YCpMep2^H348E^, YCpMep2^H194A/H348A^ or YCpMep2^H194E/H348E^. B) Growth tests, after 4 days at 29°C, on solid minimal medium containing different potassium concentrations (from 1 mM to 20 mM) and in the presence of sodium glutamate as nitrogen source. The *Saccharomyces cerevisiae* strain #228 (*mep1Δ mep2Δ mep3Δ trk1Δ trk2Δ leu2*) was transformed with the pFL38 empty plasmid (-) or with YCpMep2, YCpMep2^H194A^, YCpMep2^H194E^, YCpMep2^H348A^, YCpMep2^H348E^, YCpMep2^H194A/H348A^ or YCpMep2^H194E/H348E^.

The capacity of the ScMep2 variants to transport K^+^ was assessed in the triple-*mep*Δ *trk1*Δ *trk2*Δ strain in the presence of limiting concentrations of K^+^ (Figure 6B). Contrary to native ScMep2, and similarly to what is observed with the AmtB Twin-His variants, expression of any of the ScMep2 Twin-His variants rescues cell growth in the presence of low K^+^ concentrations, with growth efficiency depending on the variant. These data show that ScMep2 Twin-His variants can translocate K^+^, with variants possessing the H348A or H348E mutation being the most efficient in K^+^ transfer (Figure 6B). These results indicate that, as for AmtB, the hydrophobicity of the central pore of ScMep2 is involved in the transport selectivity.

### ScMep2 variants possess different transport mechanisms

It is of note that the medium used to test the ammonium transport capacity of the AmtB and ScMep2 variants contains about 180 mM K^+^. Thus, it appears that single ScMep2 His-variants are able to ensure growth in the presence of 1 mM ammonium despite a very high K^+^ concentration, indicating that potassium does not efficiently compete with ammonium recognition and transport in these variants. This indicates that the single His substitutions allow the co-existence of two transport mechanisms, as proposed for AmtB^H168D^ and AmtB^H168E^. The finding suggests that the first mechanism allows the selective transport of NH_3_ after NH_4_^+^ recognition and deprotonation as in native Mep2 and is not sensitive to high K^+^. The second mechanism, created by the substitution, could act as an ion channel pathway allowing the passage of K^+^, and likely NH_4_^+^, and would thus be sensitive to competition between both ions. This prompted us to test if the absence of growth in low ammonium conditions of cells expressing ScMep2^H194A/H348A^ and ScMep2^H194E/H348E^ could be linked to high concentrations of potassium in the medium used. We performed growth tests with a triple-*mep*Δ strain further lacking its endogenous high-affinity potassium transporter Trk1/2 in a medium allowing to simultaneously address the capacity of the variants to transport K^+^ and ammonium. As shown in Figure 7, ScMep2^H194E/H348E^ allows cell growth in the presence of 3 mM ammonium and equal or lower K^+^ concentrations (1 mM and 3 mM), indicating that the ammonium transport mechanism of this variant is highly sensitive to the potassium concentration.

**Figure 7:**
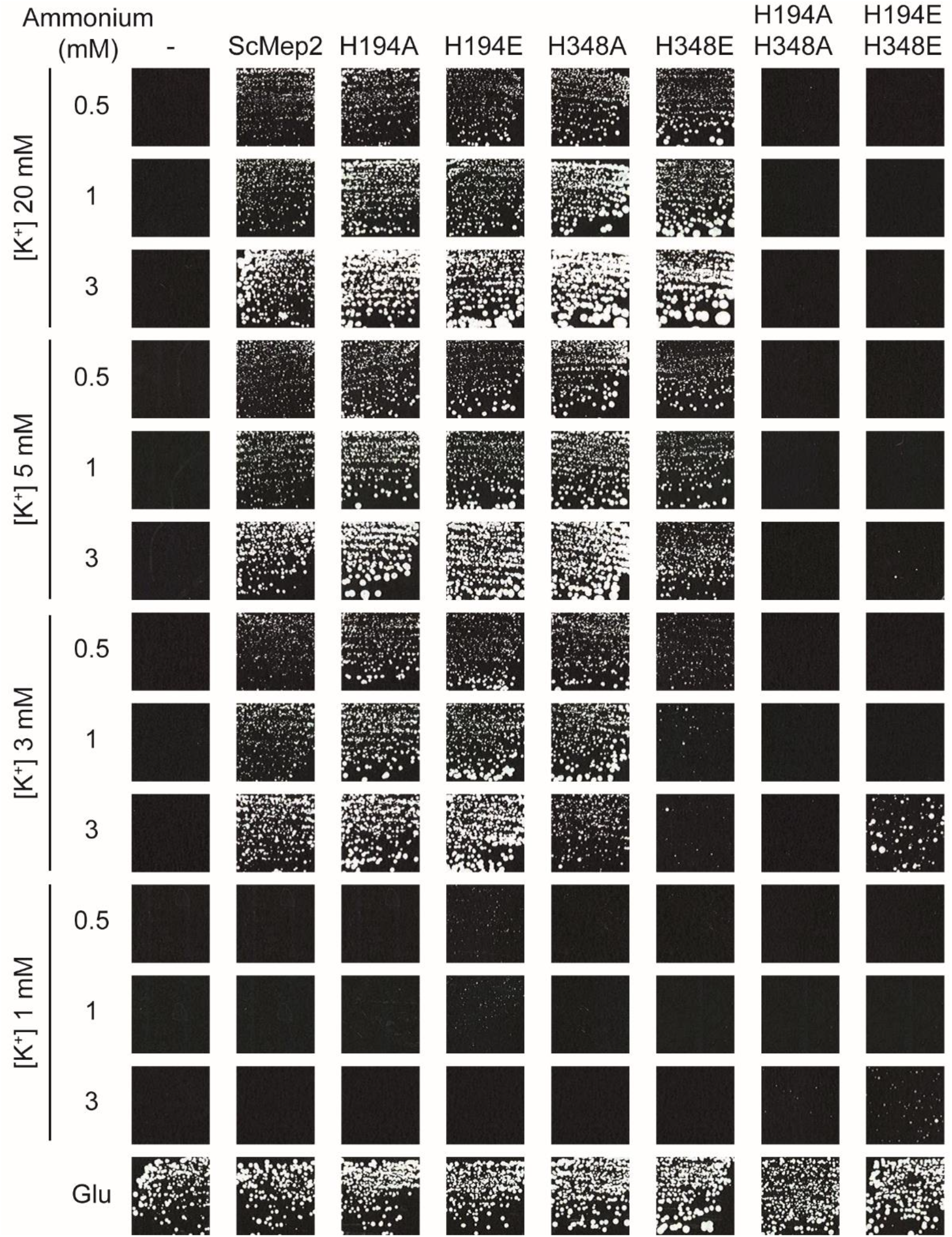
Different transport mechanisms in the ScMep2 Twin-His variants. Growth tests, after 12 days at 29°C, on solid minimal medium containing different potassium concentrations (from 1 mM to 20 mM) and in the presence of different ammonium concentrations (0.5, 1 or 3 mM) as nitrogen source. The *Saccharomyces cerevisiae* strain #228 (*mep1Δ mep2Δ mep3Δ trk1Δ trk2Δ leu2*) was double transformed with the pFL46 and pFL38 empty plasmids (-) or with pFL46 and YCpMep2, YCpMep2^H194A^, YCpMep2^H194E^, YCpMep2^H348A^, YCpMep2^H348E^, YCpMep2^H194A/H348A^ or YCpMep2^H194E/H348E^. The same solid medium containing 20 mM of potassium and potassium glutamate, as sole nitrogen source, was used as positive growth control (lower panel, the growth corresponds to day 5 at 29°C).

Consistent with the conclusions made for AmtB, these data support that simultaneous substitution of the two conserved histidines in Mep2 results in the formation of a K^+^/NH_4_^+^ channel pathway, while the transport mechanism dependent on NH_4_^+^ deprotonation is abolished. The observation that single histidine variants are able to ensure growth on low ammonium concentrations even in the presence of high potassium concentrations supports the hypothesis that both pathways, one based on ammonium deprotonation and the other linked to direct NH_4_^+^/K^+^ transport, coexist in these variants.

### Influence of the transport mechanism on the capacity of ScMep2 to induce fungal filamentation

Mep2, but not Mep1 or Mep3, is required for the dimorphic switch leading to yeast filamentation in the presence of very low ammonium concentrations ^9^. We assessed the effect of the mechanistic switch from a transporter-like to a channel-like activity on the capacity of Mep2 to induce filamentation on the appropriate SLAD medium (100 µM ammonium), as well as on SHAD medium to check for potential constitutive induction of filamentation (Figure 8A). In parallel, the ammonium transport functionality of the variants was also evaluated by growth tests on these media (Figure 8B). We show that, contrary to native ScMep2, filamentation is not observed in the presence of any of the His variants (Figure 8A). However, as Mep2^H194A/H348A^ and Mep2^H194E/H348E^ are not able to ensure growth on SLAD, probably due to the high potassium concentration (60 mM) compared to the reduced ammonium concentration (100 µM) in this medium, we cannot draw conclusions on their capacity to induce filamentation. This prompted us to test the capacity of the Twin-His variants to induce filamentation in a medium containing low potassium concentrations. Filamentation and growth tests were performed in a home-made medium, equivalent to the YNB medium and allowing to control potassium and ammonium concentrations (0.1 or 1 mM ammonium and potassium concentration ranging from 0.1 to 20 mM) (Figure 9). Again, no filamentation is observed with the Mep2 His variants (Figure 9B). However, as shown in Figure 9A, ScMep2^H194A/H348A^ and Mep2^H194E/H348E^ are not able to sustain growth at 0.1 or 1 mM ammonium, even if the potassium concentration is reduced, suggesting that ammonium transport is strongly inhibited by potassium and not sufficient to ensure growth. Of note, filamentous growth is not observed with cells expressing Mep2 at very low potassium concentrations (0.1 mM, Figure 9B). However, growth is also strongly impaired in this condition, indicating that a minimal potassium concentration is required to ensure optimal growth. Mep2-dependent filamentation is also inhibited by increasing the potassium concentration. These data suggest that altering the transport mechanism of Mep2 through Twin-His mutations impairs the capacity of the protein to induce filamentation. This result strongly supports the hypothesis that the capacity of Mep2 to induce filamentation is associated to its transport mechanism ^24,25,32,33^.

**Figure 8:**
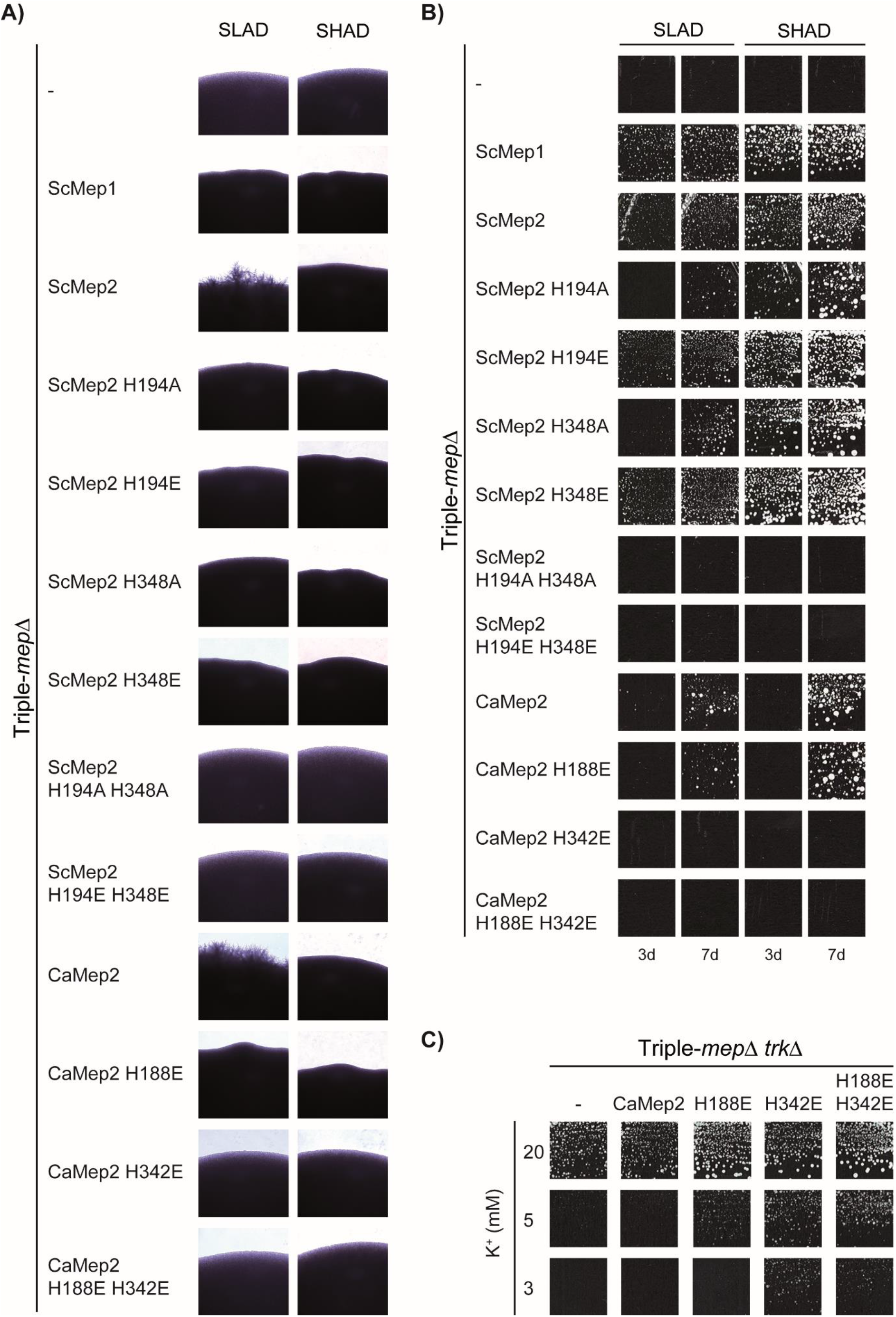
Effect of Twin-His substitutions on the capacity of ScMep2 and CaMep2 to induce fungal filamentation. Homozygous diploid triple-*mepΔ* cells (strain ZAM38) were transformed with the pFL38 empty plasmid (-) or with YCpMep2, YCpMep2^H194A^, YCpMep2^H194E^, YCpMep2^H348A^, YCpMep2^H348E^, YCpMep2^H194A/H348A^, YCpMep2^H194E/H348E^, YCpCaMep2, YCpCaMep2^H188E^, YCpCaMep2^H342E^ or YCpCaMep2^H188E/H342E^. A) Pseudohyphal growth tests of high-density cell suspensions dropped on SLAD and SHAD media at day 7. B) Growth tests of low-density cell suspensions on SLAD and SHAD media at day 3 (3d) and 7 (7d), at 29°C. C) Growth tests, after 4 days at 29°C, on solid minimal medium containing different potassium concentrations (from 2 mM to 20 mM) and sodium glutamate as nitrogen source. The *Saccharomyces cerevisiae* strain #228 (*mep1Δ mep2Δ mep3Δ trk1Δ trk2Δ leu2*) was transformed with the pFL38 empty plasmid (-) or with YCpCaMep2, YCpCaMep2^H188E^, YCpCaMep2^H342E^ or YCpCaMep2^H188E/H342E^.

**Figure 9:**
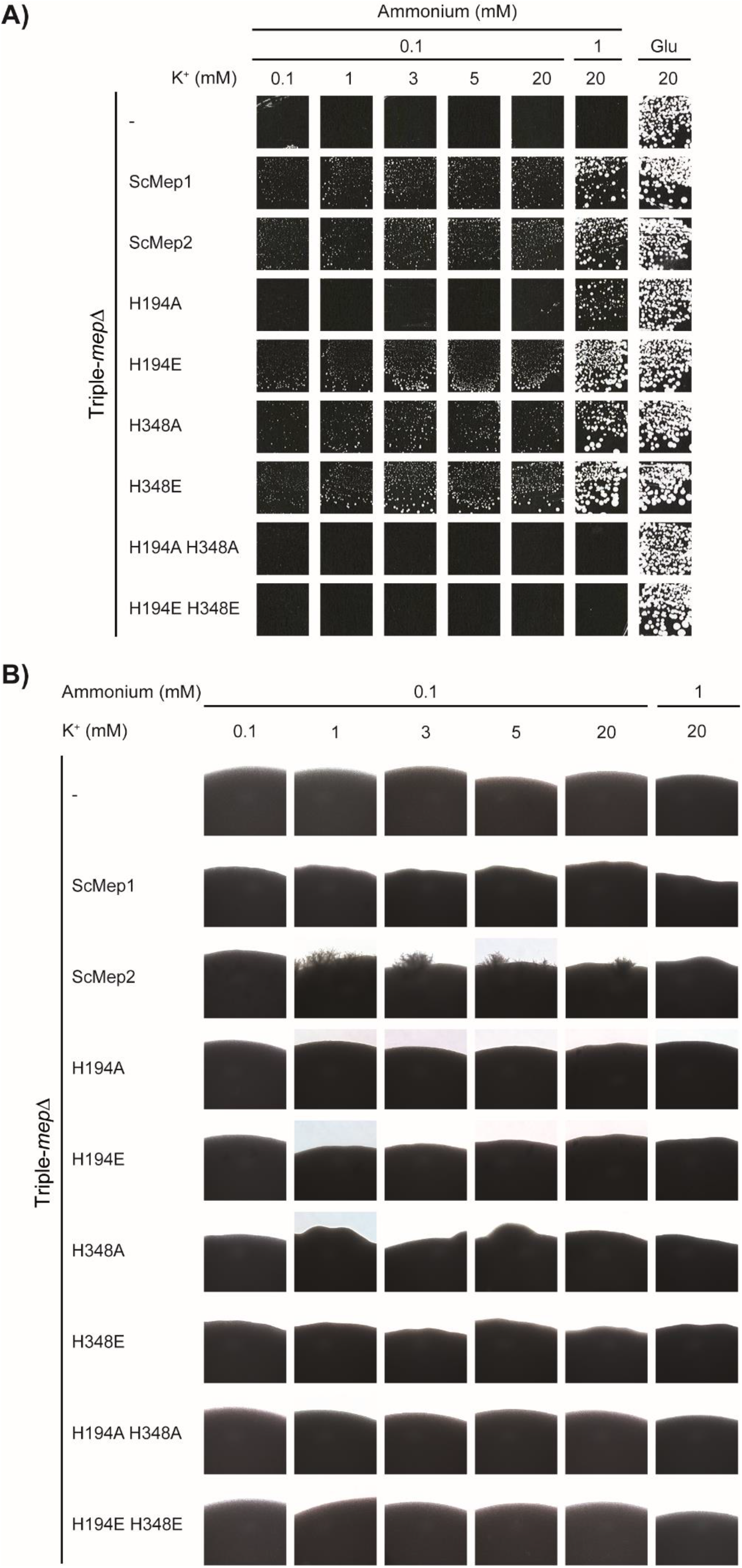
Effect of Twin-His substitutions on the capacity of ScMep2 to induce fungal filamentation at different K^+^ concentrations. Homozygous diploid triple-*mepΔ* cells (strain ZAM38) were transformed with the pFL38 empty plasmid (-) or with YCpMep1, YCpMep2, YCpMep2^H194A^, YCpMep2^H194E^, YCpMep2^H348A^, YCpMep2^H348E^, YCpMep2^H194A/H348A^ or YCpMep2^H194E/H348E^. A) Growth tests, after 7 days at 29°C, on 183 medium containing different potassium concentrations (0.1 mM - 20 mM) and 0.1 mM or 1 mM ammonium, as nitrogen source. The same medium containing 20 mM potassium and sodium glutamate (Glu), as sole nitrogen source, was used as positive growth control. B) Pseudohyphal growth tests of high-density cell suspensions dropped on 183 medium containing different potassium concentrations (from 0.1 mM to 20 mM) and 0.1 mM or 1 mM ammonium, as nitrogen source. Cells were incubated 7 days at 29°C.

### Existence of different transport mechanisms in the CaMep2 variants and influence on filamentation

CaMep2 from *Candida albicans*, the orthologue of *S. cerevisiae* Mep2, is also required to induce filamentation in this human pathogenic fungus ^34^. Expression of CaMep2 in *S. cerevisiae* cells deprived of endogenous *ScMEP2* restores filamentation, suggesting that a similar mechanism leads to filamentation in both species ^25,34^. Here, we tested the impact of Twin-His substitutions in CaMep2 by expressing the variants in *S. cerevisiae*. As observed with ScMep2, CaMep2 variants mutated in the first and/or second histidine CaMep2^H188E^, CaMep2^H342E^ and CaMep2^H188E/H342E^ are able to translocate K^+^ (Figure 8C). Growth tests on SLAD and SHAD reveal that native CaMep2 and CaMep2^H188E^ are less functional in ammonium transport than ScMep2, while CaMep2^H342E^ and CaMep2^H188E/H342E^ are completely non-functional, at least in the presence of high potassium concentrations (Figure 8B). None of the CaMep2 variants are able to induce filamentation (Figure 8A). Hence, as for ScMep2, the capacity of CaMep2 to induce filamentation also appears to be associated to its transport mechanism.

## Conclusions

The data presented show that substitutions within the Twin-His motif of Amt-Mep decrease the specificity of transport. Whilst this was previously proposed in *E. coli, S. cerevisiae* and *A. thaliana* ^22,24,35^, it was then assumed that the Twin-His motif was central for the selectivity of the transporters. In this context, the natural occurrence of a glutamic acid in the place of the first histidine in all fungal Mep1-3 proteins was unclear. Here, our findings lead us to propose a new model that explains the loss of specificity associated with variations in the Twin-His motif. We show that the modification of the Twin-His motif is associated with a change in the pore hydrophobicity increasing its hydration pattern. These findings are supported by our previous X-ray structural analysis where we showed that the introduction of a charged residue at the Twin-His position drastically enhanced the pore hydration level ^23^. This increase in hydration governs a switch of the translocation mechanism from a specific transporter, based on substrate fragmentation, to an unspecific channel-like activity, where NH_4_^+^ is transported in its hydrated form and ions of similar size are able to compete. The mechanism of substrate transport alone ensures the high specificity of the transport. This is reminiscent of the formate/nitrite transporters that ensure transport selectivity by neutralising the formate anion by protonation (deprotonation in Amt-Mep), followed by the passage of the neutral substrate through a lipophilic constriction zone ^30^. It is now essential to investigate whether the selective mechanism of Amt-Mep is conserved amongst Rhesus proteins, as RhAG mutations are associated with overhydrated stomatocytosis (OhSt), a haemolytic anaemia, which is characterized by the loss of specificity and leakage of important monovalent cations (K^+^, Na^+^) inside red blood cells ^36^.

Our results extend beyond the characterisation of a new mechanism for selectivity and offer a gateway to better understand fungal pathogenesis. The mechanism of Mep2-mediated signalling in yeast filamentation represents a long-standing debate and remains largely unsolved. Two hypotheses were raised concerning the molecular mechanism of Mep2-mediated signalling. The first and initial one is that Mep2 is a sensor, potentially interacting with signalling partners leading to the induction of filamentation ^9^. The other one postulates that the specific transport mechanism of Mep2 could mediate the signal leading to filamentation {Brito, 2020#6214. We showed that, in the Mep2 single Twin-His variants, a NH_4_^+^ deprotonation-dependent transport mechanism and a channel-like mechanism (direct NH_4_^+^ transport) coexist. We further showed that this mechanistic alteration impairs the capacity of the *S. cerevisiae* and *C. albicans* Mep2 protein to induce filamentation. Altogether, these results strongly support the hypothesis of the mechanism-mediated signalling in yeast filamentation. Filamentation is often related to the virulence of pathogenic fungi, such as the human pathogens *Candida albicans* ^*37*^, *Histoplasma capsulatum* ^38^, or *Cryptococcus neoformans* ^39^. In this context, our results are of particular importance as the characterisation of the conditions regulating the yeast dimorphism may therefore be crucial to better understand fungal virulence.

## Acknowledgements

We thank Pascale Van Vooren for support and discussion. Special thanks to Prof. Iain Hunter (Strathclyde Institute of Pharmacy and Biomedical Sciences) for invaluable discussions and help during this project.

## Materials and Methods

### Plasmids and mutageneses

Plasmids used are listed in Table S1. AmtB mutants were generated using the Quikchange XL site-directed mutagenesis kit (Agilent Technologies), following the manufacturer’s instructions. The *amtB* gene cloned into pET22b(+) was used as the template, as previously described ^11^.

Site-directed mutageneses of *ScMEP2* were performed by GeneCust, using as template YCpMep2. The *CaMEP2* (orf19.5672) gene and the mutated *CaMEP2*^*H188E*^ and *CaMEP2*^*H342E*^ genes, with the *ScMEP2* promoter (−661 to -1) and terminator (1 to 262), were synthesized and cloned in pFL38 by GeneCust. Plasmid extraction from bacterial cells was performed using the GeneJET Plasmid Miniprep Kit (Thermo Fischer). All constructs were verified by sequencing.

### AmtB expression and purification

AmtB(His)_6_ cloned into the pET22b(+) vector was overexpressed in the C43 ^40^ strain of *E. coli*, as previously describe ^11^, with minor modifications. 0.03% of *n*-dodecyl-β-d-maltoside (DDM) was used instead of 0.09% *N,N*-dimethyldodecylamine-*N*-oxide (LDAO) in the immobilised metal affinity chromatography (IMAC) and size-exclusion chromatography (SEC) buffers. AmtB was kept in the SEC buffer at 4°C prior to insertion into proteoliposomes. All the plamids used in this study are listed Table S1. All constructs were verified by sequencing.

### Insertion of AmtB into liposomes

AmtB variants were inserted into liposomes containing *E. coli* polar lipids/phosphatidylcholine (POPC) 2/1(wt/wt) as previously described ^19^. For each AmtB variant, proteoliposomes were prepared at lipid-to-protein ratios (LPRs) of 5, 10, and 50 (wt/wt). The size distribution of proteoliposomes was measured by dynamic light scattering (DLS) using a Zetasizer Nano ZS (Malvern Instruments). This analysis showed that the proteoliposomes had an average diameter of 110 nm (Figure S1). Proteoliposomes were divided into 100 µL aliquots and stored at −80°C.

To ensure that all AmtB variants were correctly inserted into the proteoliposomes, the proteoliposomes were solubilized in 2% DDM and the proteins analyzed by size exclusion chromatography using a superdex 200 (10 × 300) enhanced column. The elution profile of all variants and the wild-type were identical, showing a single monodisperse peak eluting between 10.4–10.6 ml (Figure S2). This demonstrated that all proteins were correctly folded, as trimers, in the proteoliposomes.

### Solid supported membrane electrophysiology

3 mm gold plated sensors (Nanion Technologies) were prepared according to the manufacturer’s instructions, as described previously ^41^. Proteoliposomes/empty liposomes were defrosted and sonicated in a sonication bath at 35 W for 10 seconds and diluted 10-fold in non-activating (NA) solution (Table S2), and 10 µL was added to the surface of the solid-supported membrane (SSM) on the sensor. Sensors were centrifuged at 2500 *g* for 30 minutes and stored at 4°C for a maximum of 48 hours before electrophysiological measurements. For D_2_O experiments, all the solutions were prepared using D_2_O instead of water.

All measurements were made at room temperature (21° C) using a SURFE^2^R N1 apparatus (Nanion Technologies) with default parameters ^41^. Unless otherwise stated, all measurements were carried out using pH 7 buffers. Prior to any measurements, the quality of the sensors was determined by measuring capacitance (15-30 nF) and conductance (<5 nS) and comparing to reference values provided by the manufacturer.

For functional measurements at a fixed pH, a single solution exchange protocol was used with each phase lasting 1 s ^41^. First, non-activating (NA) solution was injected onto the sensor, followed by activating (A) solution containing the substrate at the desired concentration and finally NA solution (Table S2).

Kinetic parameters were calculated using Graphpad Prism 6 and fitted according to the Michaelis-Menten equation. The decay time of the transient current was calculated by fitting the raw transient current between the apex of the peak and the baseline (after transport) with a non-linear regression using OriginPro 2017 (OriginLab). The regression was done using a one-phase exponential decay function with time constant parameter (equation below) and fit using the Levenberg Marquardt iteration algorithm.

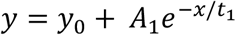

where *x*and *y* represent coordinates on the respective axis, *y0* represents the offset at a given point, *A* represents the amplitude, and *t* is the time constant.

### Yeast strains and growth conditions

The *S. cerevisiae* strains used in this study are the 31019b strain (*mep1*Δ *mep2*Δ *mep3*Δ *ura3*) ^42^, the #228 strain (*mep1*Δ *mep2*Δ *mep3*Δ *trk1*Δ *trk2*Δ *leu2 ura3*) ^43^ and the ZAM38 strain (*mep1Δ/mep1Δ mep2Δ/mep2Δ mep3Δ/mep3Δ ura3/ura3*) ^44^. Cells were grown at 29 °C. Cell transformation was performed as described previously ^45^. For growth tests on limiting ammonium concentrations, yeast cells were grown on minimal buffered (pH 6.1) medium (167) containing potassium salts ^46^. For growth tests on limiting potassium concentrations, a similar minimal buffered (pH 6.1) medium (173) where potassium salts were replaced by sodium salts was used and KCl was added at the specified concentration. 3% glucose was used as the carbon source. Nitrogen sources were added as required by the experiment and as specified in the text. The nitrogen sources used were 0.1% potassium glutamate, 0.1% sodium glutamate, or (NH_4_)_2_SO_4_ at the specified concentrations referring to the ammonium moiety.

Pseudohyphal growth tests were performed as previously described ^47^. A suspension of diploid cells was patched onto Synthetic Low Ammonium Dextrose (SLAD) and Synthetic High Ammonium Dextrose (SHAD) (0.68 % Yeast Nitrogen Base without amino acids and without (NH_4_)_2_SO_4_, containing 3 % glucose, 1 % bacteriological agar (Oxoid)), respectively supplemented with 50 µM or 0.5 mM (NH_4_)_2_SO_4_. Pseudohyphal and growth tests on limiting potassium concentrations were performed on a home-made medium (183) equivalent to Yeast Nitrogen Base medium without amino acids, (NH_4_)_2_SO_4_ and potassium salts, and containing low NaH_2_PO_4_ concentrations. (NH_4_)_2_SO_4_ and KCl were added as required by the experiment and as specified in the text. For growth tests, diploid cells were streaked on SLAD, SHAD and 183 media to follow the formation of colonies.

All growth experiments were repeated at least twice.

### Photomicroscopy

Pictures of yeast colonies were taken directly from Petri plates using a Zeiss Axio Observer Z1 microscope, driven by MetaMorph (MDS Analytical Technologies), with a 10x primary objective and a 2.5x camera adaptor.

### Molecular Dynamics Simulations

The AmtB trimer (PDB code: 1U7G) ^10^ was processed using the CHARMM-GUI web server ^48^. Any mutations inserted during the crystallization process were reverted to the wild-type form. The studied mutations were introduced into the protein using Pymol. The N-termini and C-termini of the subunits were capped with acetyl and N-methyl amide moieties, respectively. The protein was then inserted into a membrane patch of *xy*-dimensions 13 × 13 nm. Unless otherwise specified, a membrane composition of palmitoyl oleoyl phosphatidyl ethanolamine and palmitoyl oleoyl phosphatidyl glycine (POPE/POPG) at a 3:1 ratio was used in order to approximate the composition of a bacterial cytoplasmic membrane. We employed the CHARMM36 forcefield for the protein and counter ions ^49^.The water molecules were modeled with the TIP3P model ^50^. Water bonds and distances were constrained by the Settle method ^51^, and bonds involving hydrogen by the LINCS method ^52^. In simulations without ammonium, K^+^ and Cl^-^ions were added to neutralize the system and obtain a bulk ionic concentration of 250 mM. In simulations with ammonium, K^+^ was replaced by NH_4_^+^.

After a steepest descent energy minimization, the system was equilibrated by six consecutive equilibration steps using position restraints on heavy atoms of 1000 kJ/mol.nm^2^. The first three equilibration steps were conducted in an NVT ensemble, applying a Berendsen thermostat ^53^ to keep the temperature at 310K. The subsequent steps were conducted under an NPT ensemble, using a Berendsen barostat ^53^ to keep the pressure at 1 bar. Production molecular dynamics simulations were carried out using a v-rescale thermostat ^54^ with a time constant of 0.2 ps, and a Berendsen barostat with semi-isotropic coupling. A timestep of 2 fs was used throughout the simulations.

**Figure S1:**
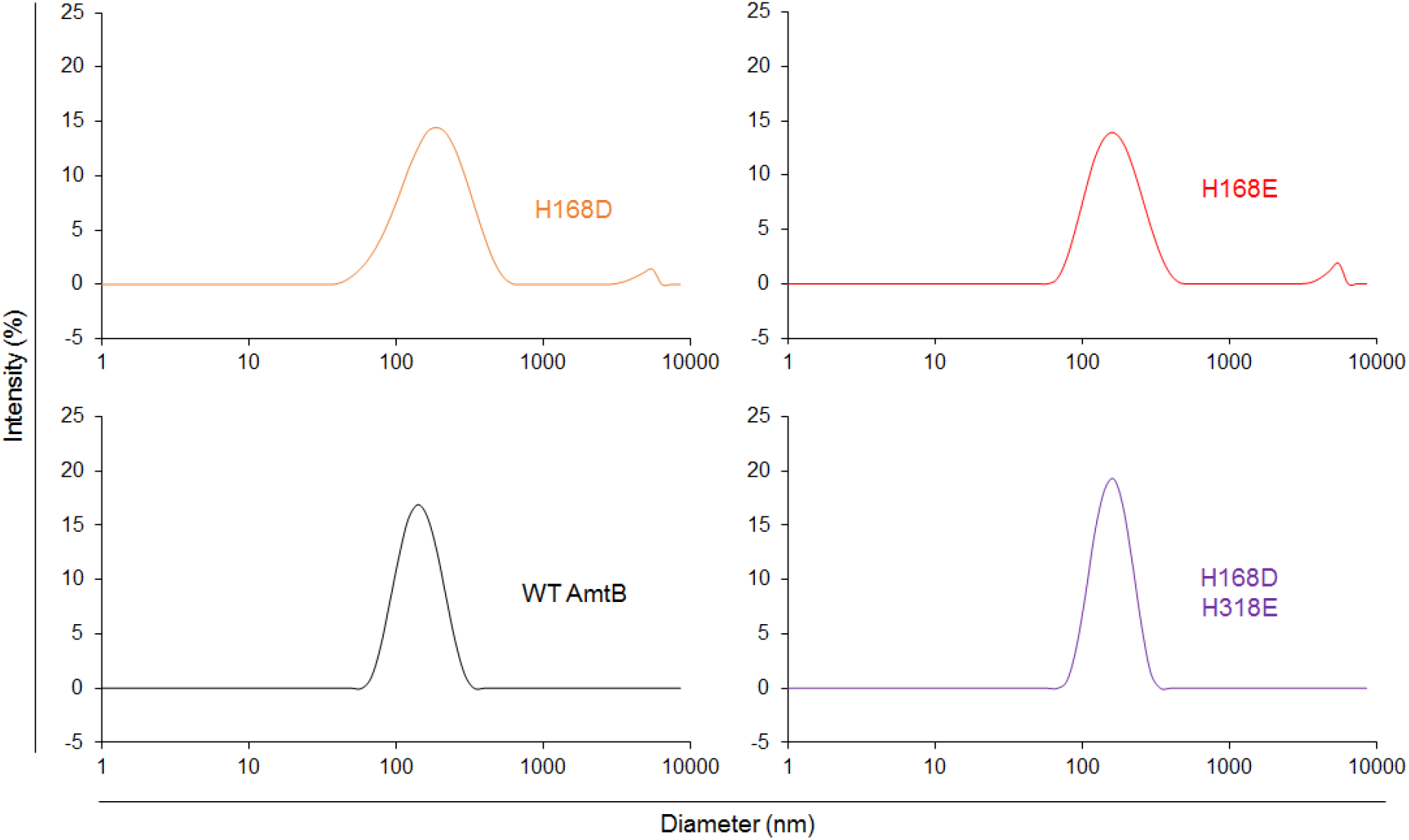
Size distribution of the proteoliposomes containing wild-type and variants of AmtB. Dynamic light scattering was used to determine the number-weighted distribution of liposome sizes in the detection reagent. The distribution was monodisperse, with a mean diameter of 110 nm.

**Figure S2:**
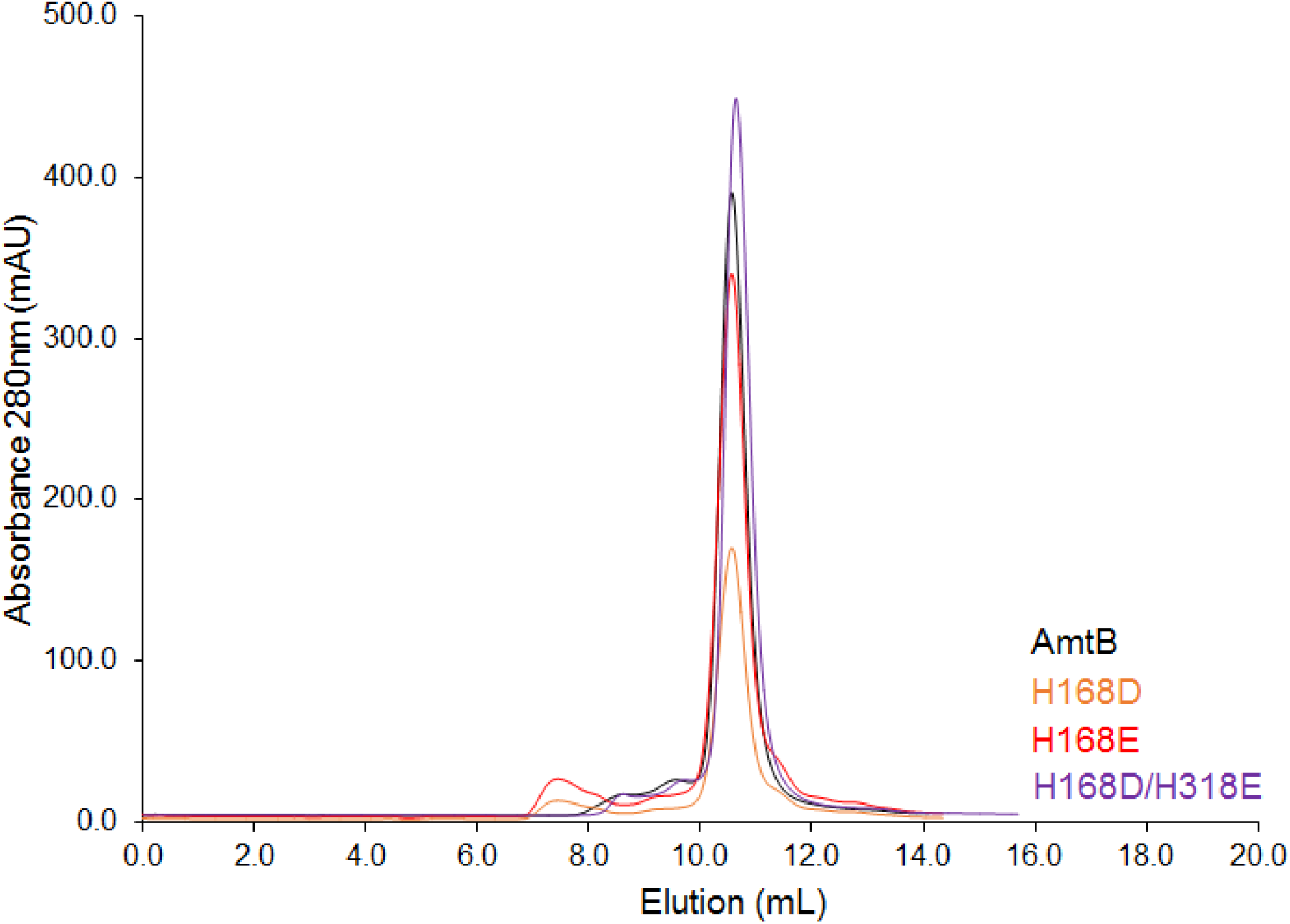
Size exclusion chromatography analysis of AmtB variants. Gel filtration trace (Superdex 200 10/300 increase) of wild-type AmtB and variants after solubilization of the proteoliposome in 2% DDM. All of the AmtB variants elute at ∼11.5 ml.

**Table S1:** Plasmids used in this study.

| Plasmid | Description | Reference |
| --- | --- | --- |
| ***E. coli*** |  |  |
| pET22b (+) | High copy bacterial expression vector | Novagen |
| pZheng | pET22b-AmtB(His)<sub>6</sub> | 11 |
| pAJ2039 | pET22b-AmtB(His)<sub>6</sub><sup>H168D</sup> | 23 |
| pGW1 | pET22b-AmtB(His)<sub>6</sub><sup>H168E</sup> | This study |
| pGW2 | pET22b-AmtB<sup>H168D/H318E</sup> | This study |
| ***S. cerevisiae*** |  |  |
| pFL38 | *CEN-ARS URA3* | 55 |
| pFL46 | *2μ LEU2* | 55 |
| pDR195 | *2μ URA3* | 56 |
| pGW6 | pDR195-*AMTB*<sup>H168E</sup> | This study |
| pGDM8 | pDR195-*AMTB*<sup>H168D</sup> | This study |
| pGDM9 | pDR195-*AMTB*<sup>H168D/H318E</sup> | This study |
| YCpMep2 | YCpFL38 *MEP2* | 42 |
| YCpMep1 | YCpFL38 *MEP1* | 5 |
| YCpMep2<sup>H194E</sup> | YCpFL38 *MEP2*<sup>H194E</sup> | 24 |
| YCpMep2<sup>H194A</sup> | YCpFL38 *MEP2*<sup>H194A</sup> | This study |
| YCpMep2<sup>H348E</sup> | YCpFL38 *MEP2*<sup>H348E</sup> | This study |
| YCpMep2<sup>H348A</sup> | YCpFL38 *MEP2*<sup>H348A</sup> | 24 |
| YCpMep2<sup>H194A, H348A</sup> | YCpFL38 *MEP2*<sup>H194A, H348A</sup> | This study |
| YCpMep2<sup>H194E, H348E</sup> | YCpFL38 *MEP2*<sup>H194E, H348E</sup> | This study |
| YCpCaMep2 | YCpFL38 PROM *ScMEP2*-*CaMEP2*-TERM *ScMEP2* | This study |
| YCpCaMep2<sup>H188E</sup> | YCpFL38 PROM<br>*ScMEP2*-<br>*CaMEP2*<sup>H188E</sup>-TERM<br>*ScMEP2* | This study |
| YCpCaMep2<sup>H342E</sup> | YCpFL38 PROM<br>*ScMEP2*-<br>*CaMEP2*<sup>H348E</sup>-TERM<br>*ScMEP2* | This study |
| YCpCaMep2<sup>H188E, H342E</sup> | YCpFL38 PROM<br>*ScMEP2*-<br>*CaMEP2*<sup>H188E, H348E</sup>-<br>TERM *ScMEP2* | This study |

**Table S2:**
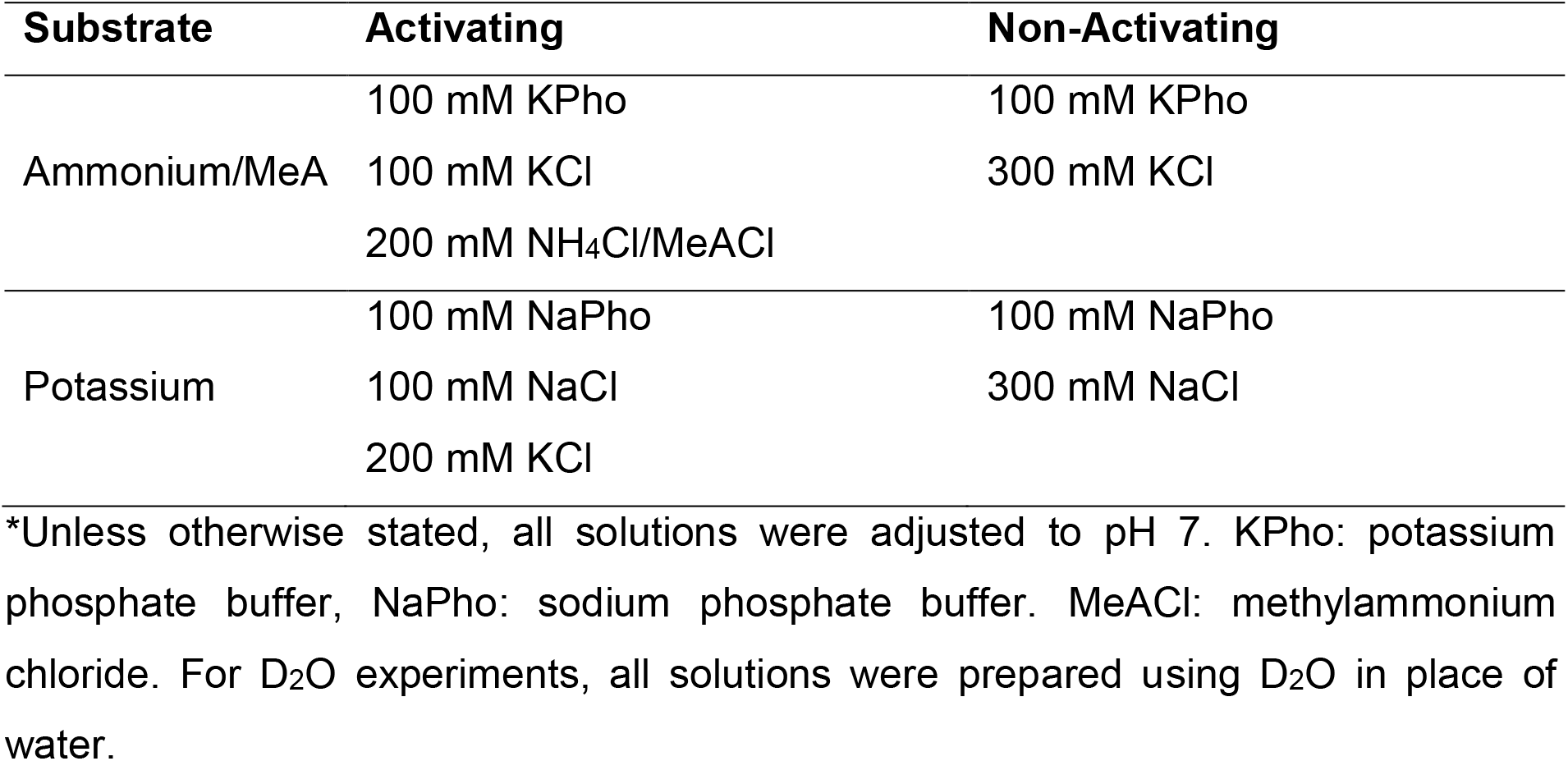
Solid-supported membrane electrophysiology solutions.

## References

1 von Wiren, N. & Merrick, M. Regulation and function of ammonium carriers in bacteria, fungi and plants. Trends Genet. 9, 95–120 (2004).

2 Martinelle, K., Westlund, A. & Haggstrom, L. Ammonium ion transport-a cause of cell death. Cytotechnology 22, 251–254 (1996).

3 Posset, R. et al. Age at disease onset and peak ammonium level rather than interventional variables predict the neurological outcome in urea cycle disorders. J Inherit Metab Dis 39, 661–672 (2016).

4 Huang, C. H. & Peng, J. Evolutionary conservation and diversification of Rh family genes and proteins. Proc. Natl. Acad. Sci. U.S.A 102, 15512–15517 (2005).

5 Marini, A. M., Vissers, S., Urrestarazu, A. & Andre, B. Cloning and expression of the MEP1 gene encoding an ammonium transporter in Saccharomyces cerevisiae EMBO J. 13, 3456–3463 (1994).

6 Ninnemann, O., Jauniaux, J. C. & Frommer, W. B. Identification of a high affinity NH4+ transporter from plants. EMBO J. 13, 3464–3471 (1994).

7 Marini, A. M., Urrestarazu, A., Beauwens, R. & Andre, B. The Rh (Rhesus) blood group polypeptides are related to NH4+ transporters. Trends Biochem. Sci. 22, 460–461 (1997).

8 Marini, A. M. et al. The human Rhesus-associated RhAG protein and a kidney homologue promote ammonium transport in yeast. Nat. Genet. 26, 341–344 (2000).

9 Lorenz, M. C. & Heitman, J. The MEP2 ammonium permease regulates pseudohyphal differentiation in Saccharomyces cerevisiae EMBO J. 17, 1236–1247 (1998).

10 Khademi, S. et al. Mechanism of ammonia transport by Amt/MEP/Rh: structure of AmtB at 1.35Å. Science 305, 1587–1594 (2004).

11 Zheng, L., Kostrewa, D., BernŠche, S., Winkler, F. K. & Li, X. D. The mechanism of ammonia transport based on the crystal structure of AmtB of E. coli. Proc. Natl. Acad. Sci. U.S.A. 101, 17090–17095 (2004).

12 Lupo, D. et al. The 1.3-Å resolution structure of Nitrosomonas europaea Rh50 and mechanistic implications for NH3 transport by Rhesus family proteins. Proc. Natl. Acad. Sci. U.S.A 104, 19303–19309 (2007).

13 Gruswitz, F. et al. Function of human Rh based on structure of RhCG at 2.1 Å. Proc. Natl. Acad. Sci. U.S.A 107, 9638–9643 (2010).

14 van den Berg, B. et al. Structural basis for Mep2 ammonium transceptor activation by phosphorylation. Nat Commun 7, 11337 (2016).

15 Andrade, S. L., Dickmanns, A., Ficner, R. & Einsle, O. Crystal structure of the archaeal ammonium transporter Amt-1 from Archaeoglobus fulgidus Proc. Natl. Acad. Sci. U.S.A 102, 14994–14999 (2005).

16 Ariz, I. et al. Nitrogen isotope signature evidences ammonium deprotonation as a common transport mechanism for the AMT-Mep-Rh protein superfamily. Sci Adv 4, eaar3599. doi:10.1126/sciadv.aar3599 (2018).

17 Dias Mirandela, G. et al. Merging In-Solution X-ray and Neutron Scattering Data Allows Fine Structural Analysis of Membrane-Protein Detergent Complexes. J. Phys. Chem. Lett. 9, 3910–3914 (2018).

18 Wacker, T., Garcia-Celma, J. J., Lewe, P. & Andrade, S. L. Direct observation of electrogenic NH4(+) transport in ammonium transport (Amt) proteins. Proc. Natl. Acad. Sci. U.S.A 111, 9995–10000 (2014).

19 Mirandela, G. D., Tamburrino, G., Hoskisson, P. A., Zachariae, U. & Javelle, A. The lipid environment determines the activity of the Escherichia coli ammonium transporter AmtB. FASEB J, 33, 1989–1999 (2018).

20 Javelle, A. et al. Structural and mechanistic aspects of Amt/Rh proteins. J. Struct. Biol. 158, 472–481 (2007).

21 Merrick, M. et al. The Escherichia coli AmtB protein as a model system for understanding ammonium transport by Amt and Rh proteins. Transfus. Clin. Biol. 13, 97–102 (2006).

22 Williamson, G. et al. A two-lane mechanism for selective biological ammonium transport. Elife 9, e57183 (2020).

23 Javelle, A. et al. An unusual twin-his arrangement in the pore of ammonia channels is essential for substrate conductance. J. Biol. Chem. 281, 39492–39498 (2006).

24 Boeckstaens, M., Andre, B. & Marini, A. M. Distinct transport mechanisms in yeast ammonium transport/sensor proteins of the Mep/Amt/Rh family and impact on filamentation. J. Biol. Chem. 283, 21362–21370 (2008).

25 Brito, A. S., Neuhauser, B., Wintjens, R., Marini, A. M. & Boeckstaens, M. Yeast filamentation signaling is connected to a specific substrate translocation mechanism of the Mep2 transceptor. PLoS Genet 16, e1008634, (2020).

26 Hess, D. C., Lu, W., Rabinowitz, J. D. & Botstein, D. Ammonium toxicity and potassium limitation in yeast. PLoS Biol 4, e351 (2006).

27 Loque, D., Mora, S. I., Andrade, S. L., Pantoja, O. & Frommer, W. B. Pore mutations in ammonium transporter AMT1 with increased electrogenic ammonium transport activity. J. Biol. Chem. 284, 24988–24995 (2009).

28 Wang, J. et al. Ammonium transport proteins with changes in one of the conserved pore histidines have different performance in ammonia and methylamine conduction. PLoS One 8, e62745 (2013).

29 Roon, R. J., Larimore, F. & Levy, J. S. Inhibition of amino acid transport by ammonium ion in Saccharomyces cerevisiae. J. Bacteriol. 124, 325–331 (1975).

30 Wiechert, M. & Beitz, E. Mechanism of formate-nitrite transporters by dielectric shift of substrate acidity. EMBO J. 36, 949–958 (2017).

31 Dubois, E. & Grenson, M. Methylamine/ammonia uptake systems in Sacharomyces cerevisiae: multiplicity and regulation. Mol. Gen. Genet. 175, 67–76 (1979).

32 Lorenz, M. C. & Heitman, J. Regulators of pseudohyphal differentiation in Saccharomyces cerevisiae identified through multicopy suppressor analysis in ammonium permease mutant strains. Genetics 150, 1443–1457 (1998).

33 Lorenz, M. C. & Heitman, J. Yeast pseudohyphal growth is regulated by GPA2, a G protein alpha homolog. EMBO J. 16, 7008–7018 (1997).

34 Biswas, K. & Morschhauser, J. The Mep2p ammonium permease controls nitrogen starvation-induced filamentous growth in Candida albicans Mol. Microbiol. 56, 649–669 (2005).

35 Ganz, P. et al. A twin histidine motif is the core structure for high-affinity substrate selection in plant ammonium transporters. J. Biol. Chem. 295, 3362–3370 (2020).

36 Bruce, L. J. et al. The monovalent cation leak in overhydrated stomatocytic red blood cells results from amino acid substitutions in the Rh-associated glycoprotein. Blood 113, 1350–1357 (2009).

37 Lo, H. J. et al. Nonfilamentous C. albicans mutants are avirulent. Cell 90, 939–949 (1997).

38 Maresca, B. & Kobayashi, G. S. Dimorphism in Histoplasma capsulatum: a model for the study of cell differentiation in pathogenic fungi. Microbiol. Rev. 53, 186–209 (1989).

39 Wickes, B. L., Edman, U. & Edman, J. C. The Cryptococcus neoformans STE12alpha gene: a putative Saccharomyces cerevisiae STE12 homologue that is mating type specific. Mol. Microbiol. 26, 951–960 (1997).

40 Miroux, B. & Walker, J. E. Over-production of proteins in Escherichia coli: mutant hosts that allow synthesis of some membrane proteins and globular proteins at high levels. J. Mol. Biol.260, 289–298 (1996).

41 Bazzone, A., Barthmes, M. & Fendler, K. SSM-Based Electrophysiology for Transporter Research. Methods Enzymol. 594, 31–83 (2017).

42 Marini, A. M., Soussi-Boudekou, S., Vissers, S. & Andre, B. A family of ammonium transporters in Saccharomyces cerevisae Mol. Cell. Biol. 17, 4282–4293 (1997).

43 en Hoopen, F. et al. Competition between uptake of ammonium and potassium in barley and Arabidopsis roots: molecular mechanisms and physiological consequences. J. Exp. Bot. 61, 2303–2315 (2010).

44 Marini, A. M., Boeckstaens, M., Benjelloun, F., Cherif-Zahar, B. & Andre, B. Structural involvement in substrate recognition of an essential aspartate residue conserved in Mep/Amt and Rh-type ammonium transporters. Curr. Genet. 49, 364–374 (2006).

45 Gietz, D., St Jean, A., Woods, R. A. & Schiestl, R. H. Improved method for high efficiency transformation of intact yeast cells. Nucleic. Acids Res. 20, 1425 (1992).

46 Jacobs, P., Jauniaux, J. C. & Grenson, M. A cis-dominant regulatory mutation linked to the argB-argC gene cluster in Saccharomyces cerevisiae. J. Mol. Biol. 139, 691–704 (1980).

47 Gimeno, C. J., Ljungdahl, P. O., Styles, C. A. & Fink, G. R. Unipolar cell divisions in the yeast S. cerevisiae lead to filamentous growth: regulation by starvation and RAS. Cell 68, 1077–1090 (1992).

48 Lee, J. et al. CHARMM-GUI Input Generator for NAMD, GROMACS, AMBER, OpenMM, and CHARMM/OpenMM Simulations Using the CHARMM36 Additive Force Field. J. Chem. Theory Comput. 12, 405–413 (2016).

49 Best, R. B. et al. Optimization of the additive CHARMM all-atom protein force field targeting improved sampling of the backbone phi, psi and side-chain chi(1) and chi(2) dihedral angles. J. Chem. Theory Comput. 8, 3257–3273 (2012).

50 Jorgensen, W. L., Chandrasekhar, J., Madura, J. D., Impey, R. W. & Klein, M. L. Comparison of simple potential functions for simulating liquid water. J. Chem. Physics 79, 926–935 (1983).

51 Miyamoto, S. & Kollman, P. A. Settle: An analytical version of the SHAKE and RATTLE algorithm for rigid water models. J. Comput. Chem. 13, 952–962 (1992).

52 Hess, B., Bekker, H., Berendsen, H. J. C. & Fraaije, J. G. E. M. LINCS: A linear constraint solver for molecular simulations. J. Comput. Chem. 18, 1463–1472 (1997).

53 Berendsen, H. J. C., Postma, J. P. M., van Gunsteren, W. F., DiNola, A. & Haak, J. R. Molecular dynamics with coupling to an external bath. J. Chem. Physics 81, 3684–3690 (1984).

54 Bussi, G., Donadio, D. & Parrinello, M. Canonical sampling through velocity rescaling. J. Chem. Physics 126, 014101 (2007).

55 Bonneaud, N. et al. A family of low and high copy replicative, integrative and single-stranded S. cerevisiae/E. coli shuttle vectors. Yeast 7, 609–615 (1991).

56 Rentsch, D. et al. NTR1 encodes a high affinity oligopeptide transporter in Arabidopsis FEBS Letters 370, 264–268 (1995).

